# Detection of the germline specific NAC protein paralog enriched by intrinsically disordered regions in *Drosophila melanogaster*

**DOI:** 10.1101/2021.03.09.434555

**Authors:** Galina L. Kogan, Elena A. Mikhaleva, Oxana M. Olenkina, Sergei S. Ryazansky, Oxana V. Galzitskaya, Yuri A. Abramov, Toomas A. Leinsoo, Natalia V. Akulenko, Vladimir A. Gvozdev

## Abstract

Nascent polypeptide associated complex (NAC) consisting of α- and β-subunits is an essential conserved ubiquitously expressed ribosome-associated protein in eukaryotes. NAC is considered as a chaperone and co-translational regulator of nascent protein sorting providing homeostasis of cellular proteins. Here we discovered the germinal cell specific NAC (gNAC) homologue, which differs from the ubiquitously expressed NAC by the presence of expanded intrinsically disordered regions (IDRs) at the N- and C-ends of the α- and β-subunits, respectively. We propose these evolutionary acquisition of long IDRs drive gNAC to endow both the specific conformational plasticity for binding client proteins and novel functions regulated by post-transcriptional modifications (PTM). At the same time, we demonstrated that the well-known lethal effect of the loss of ubiquitous NAC-β is suppressed by ectopic expression of its germinal paralog indicating the absence of strict functional differences between the ubiquitous and germline NAC-β subunit paralogs for protein homeostasis.

## Introduction

The evolutionally conservative NAC (nascent polypeptide associated complex consisting of α and β subunits) interacts with translating ribosomes and functions as a ubiquitous ATP-independent chaperone protein in eukaryotes [1, 2]. It is suggested that most translating ribosome are associated with NAC [3]. NAC is involved in co-translational protein folding [4] and delivery of mitochondrial precursor proteins [5]. Recently new mechanistical insights were achieved in the studies of NAC-β subunit function, it was shown that N-terminal of NAC-β subunit is able to be inserted deeply into the ribosomal tunnel, close to polypeptide transferase center, to sense nascent polypeptide [6].The sensing NAC-β ability is coupled with its flexibility and conformational switches that either activates or prevents its associations with the translocon complex of endoplasmatic reticulum (ER) or signal recognizing particle (SRP). These conformational changes of flexible NAC structure are suggested to regulate correct protein sorting for membrane components or secretion. NAC modulates protein transmission to ER lumen or cytosol, ensuring specific co-translational protein biogenesis pathway [1, 2].

Here we focus on the specific paralogs of NAC-β and NAC-α proteins expressed in the germ cells in *D.melanogaster*. Recently, for the first time to our knowledge, we have discovered the expression of tissue specific NAC in metazoa by revealing the specific NAC-β protein expression exclusively in the germ cells of *D.melanogaster* testis [7]. Here we extended this result detecting germinal NAC-α as an immediate product of the gene CG4415 in the *D.melanogaster* genome thus demonstrating the existence of the germline specific heterodimeric NAC-αβ (gNAC). The CG4415 gene protein and previously annotated representative of a set of the germinal NAC-β encoding genes [8] we found here to be participated in gNAC generation. Here we also demonstrate that both germinal α- and β-subunits have evolved to acquire extensive intrinsically disordered regions (IDRs), primarily at the N- and C-ends, respectively. The IDRs are known to be important chaperone functional regions [9, 10] and their significant increase in gNAC may indicate a complex role for IDRs in client detection of conformational switches in germ cells. The specific functions of gNAC are still mysterious, but it is believed that acquired IDRs are generally prone to extensive post-translational modifications (PTMs) [11] as well as formation of functional chaperone function in protein-protein interactions [12]. Our results also allow us to suggest that the evolutionary emergence of the specific germline NAC with extended IDRs enhances the ability of these paralogs to execute various intermolecular interactions and PTM acquisitions. This suggestion is indirectly consistent with the observed ability of gNAC-β to mediate some multifaceted NAC functions of its ubiquitously expressed counterpart, since ectopic expression of gNAC-β is shown here to be effective to rescue the known lethal effect of the loss of ubiquitous NAC-β subunit [13].

We have found earlier the association of the germinal β-subunit with ribosomes and polysomes in the testes [7]. Taking into account a wide discussion devoted to the existence of special ribosomes performing cell specific functions (for reviews [14–19]) due to selective mRNAs translation, germinal NAC-associated ribosomes can be considered as specialized ones. It has been shown that ribosome specificity is determined not only by specific ribosomal proteins [20] or ribosomal RNA distinct structural peculiarities [16], but also by ribosome associated factors [21, 22]. The discovery of tissue specific NAC opens up new perspectives in the studies of specialized ribosomes role [21] in the development of the germline.

We revealed a distinct feature of the gNAC subunits carrying extended IDRs known to be involved in protein-protein interactions as well as in increased ability to accumulate post translational modifications [23, 24]. Currently, much attention is paid to the role of IDRs as protein recruitment regions in the formation of protein complexes (hubs) that provide a number of important cellular processes, including transcription [25, 26], DNA repair [27], cell signaling [28], germ cell development [29] and more others. The ribosome-associated complexes include apart from NAC the Hsp40/70 chaperones, SRP complex and others proteins, which coordinated interactions provide a system responsible for regulation of protein homeostasis [1] using flexible IDRs.

## Results

### Detection of the germ cell specific α- and β-paralogs forming the heterodimeric germinal NAC

Previously, we have succeeded to annotate several β*NACtes* genes amplified on the X-chromosome of *D.melanogaster*. These genes are represented by two distinct closely related, but non-adjacent pairs and an additional pair outside these two pairs, as paralogs of the ubiquitously expressed *bicaudal (bic)* gene encoding β-subunit of NAC protein [30] as well as in Flybase. β*NACtes* are paralogs of the ubiquitously expressed *bicaudal (bic)* gene encoding β-subunit of NAC protein [7]. The specific expression of these genes in testis spermatocytes, but not in somatic testes cells, has been demonstrated [7]. The gene CG4415 product carries α-NAC domain and was shown to be expressed in embryonic gonad (Fisher et al., 2012, Flybase). Moreover, a network analysis of CG18313 (*NAC*-β*tes*) using the String database (v. 10.5) [31] indicated its product association with CG4415 (NAC-α subunit).To directly demonstrate the heterodimerization of the putative germinal α-NAC and β-NACtes subunits, we performed co-transfection of S2 somatic *D.melanogaster* cells with two plasmids expressing Flag-tagged NAC-β and HA-tagged NAC-α subunits. The putative heterodimer was immunoprecipitated using anti-FLAG or anti-HA beads and its subsequent WB-analysis revealed both subunit proteins into two distinct IP samples (Fig.1 A). Taking into account the detection of RNA transcripts of the CG4415 gene in embryonic primordial gonads revealed by *in situ* hybridization (Fig. 1B, Fisher et al.2012), we performed immunostaining of embryos using generated rabbit anti-βNACtes Abs (Fig.1 C, D, E, F). The similar expression patterns of both αNAC mRNA (Fisher et al. 2012, BDGP in situ homepage, fruitfly.org) and βNACtes subunits in primordial embryonic gonad (Fig_ 1B,C) allows us to further use the gNAC designation for heterodimeric germline specific NAC protein, which β-subunit has been earlier termed as βNACtes expressed exclusively in germinal cells in testis [7]. The gNAC appears to be extremely flexible as we detected expanded IDR sequences in both of its subunits compared to the ubiquitous NAC protein.

**Figure 1.**
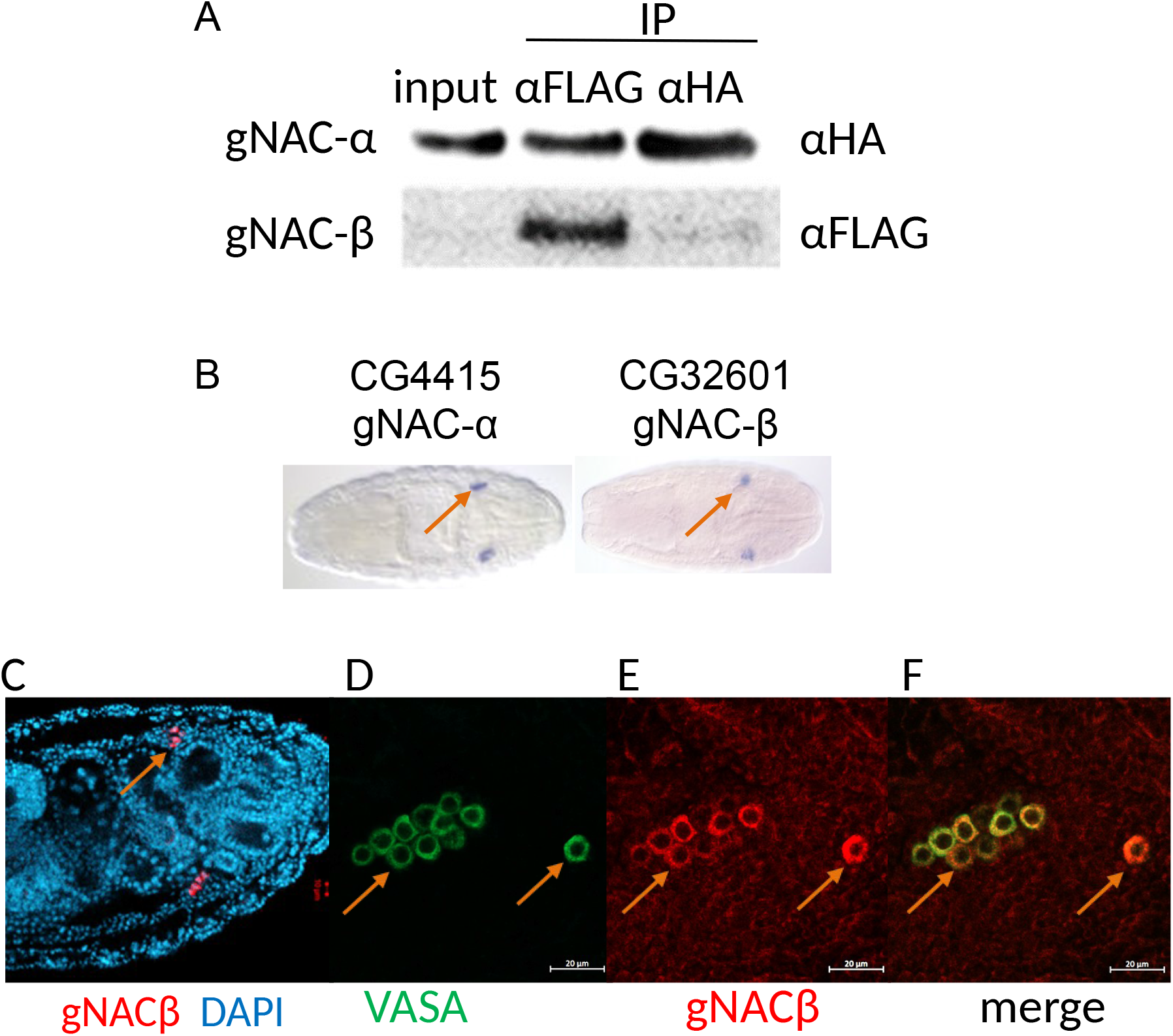
Detection of artificial gNAC-α/gNAC-β heterodimer in S2 cells and expression of gNAC-β in embryo. **A.** Generation of gNAC after S2 cells co-transfection with plasmids expressing Ac5.1 gNAC-αHA and pAc5.1-FLAG-gNAC-β subunits. **B.** FISH hybridization to detect CG4415 and CG32601 genes expression (Fisher et al., 2012); **C-F,** immunostaining of gNAC-β in 13-14th stage embryos; arrows show primordial gonads (B,C,D) or groups of migrating germ cells (E,F).

### The germinal α- and β-subunits are enriched with IDRs

The study of the conformation of the ubiquitously expressed NAC from *Caenorhabditis elegans* revealed its flexibility, which ensures its ability to bind various protein substrates, irrespective of whether they are folded or intrinsically disordered [32]. The discovery of the germline paralogs of both NAC subunits prompts us to compare the predicted conformational flexibility of their sequences with ubiquitous analogues by assessing the IDRs outside the folded NAC domain responsible for αβ dimerization. The profiles of the disordered regions were obtained using the IsUnstruct program [33, 34] (Fig. 2). The C-terminus of both ubiquitous and germinal α-subunits contain the NAC-domain and the ubiquitin-activated domain (UBA), the function of which remains enigmatic [35]. The germinal α-subunit has a threefold increase in IDR at the N-terminus in comparison with the ubiquitous analogue. The ubiquitous β-subunits carry a conserved positively charged motif at the N-end of the ribosome associated region [1] and the NAC dimerizing domain, but germinal counterpart conserving this charged motif has a more than twofold increase in IDR at the C-terminus. Thus, a significant difference in amino acid sequence patterns between the germinal and ubiquitous subunits lies in the extensions of germinal IDRs. It should be noted that the positive sequence of N-termini of the NAC-β subunit from C. *elegans* was shown recently not only to be responsible for ribosome binding, but also, as predicted, is unstructured and capable of expressing ribosome independent chaperone activity [36].

**Figure 2.**
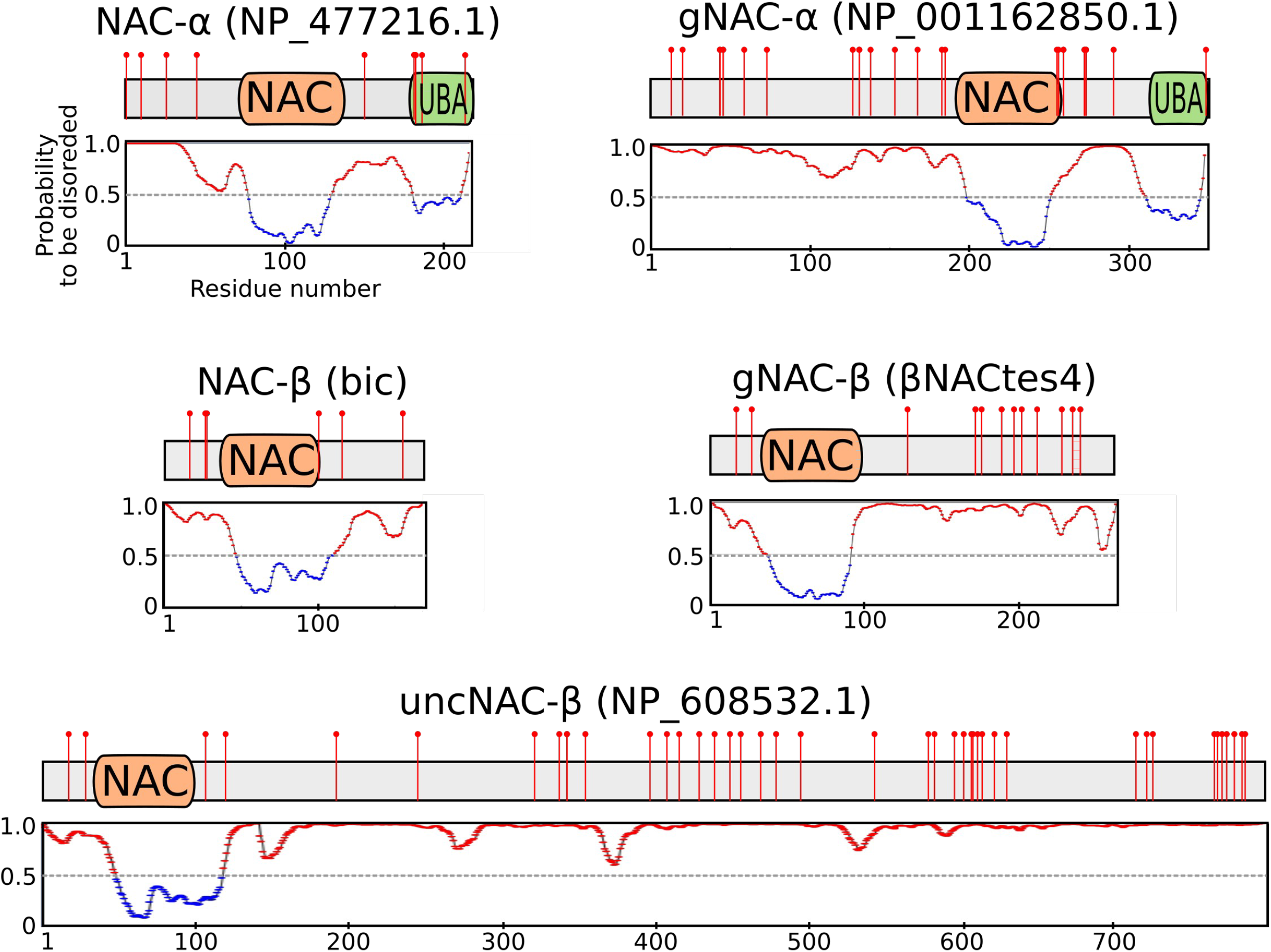
Schematic representation of NAC containing polypeptides with the plot providing the distribution of disorder propensity. The predicted posphosites are marked by red lollipops.

To get some clues about the functional significance of the discovered IDRs, and considering that phosphorylation is regarded as one of most common types of PTM in IDRs, we used ELM (eukaryotic linear motif resource for functional sites in proteins) to predict phosphosites and docking motifs for protein kinases along the gNAC-β sequences (Fig.2), while these estimations seem to be not very reliable and require an experimental confirmation. Long IDRs (more than 100 amino acids) are known to be preferentially phosphorylated [37]. Many of the predicted phospho-sites along the elongated germinal IDRs are presented in Fig.2, including an uncharacterized NAC-domain containing protein, uncNAC-β, whose phosphosite clusters correspond to disordered regions. We traced the diversity of the predicted phosphosites (Fig.2) by comparing four highly homologous copies of NAC-β gene [7], previously designated as the β-NACtes ones (Flybase.org) (Fig.S1A). The tes4 copy demonstrates the emergence of phosphosite within the newly acquired amino acid stretch and acquisition of the adjacent phosphosite. At the same time, two phosphosites were lost with the very tes4 C-terminus, which was conserved in the other three copies.

Of particular interest is the evolutionary origins of gNAC with a focus on its IDR orthologous regions. We constructed the phylogenetic tree including 69 *Drosophilidae* species whose genomic assemblies were downloaded from the NCBI. The assemblies were analyzed by BUSCO according to [38] to identify universal single copy orthologs after multiple alignment. The generated phylogenic tree is in a good agreement with others previously reported (Fig 3) (e.g. «Drosophila 25 Species Phylogeny» 2017 Dataset posted on 28.09.2017). We found that the genomes of all mentioned species from the *Sophophora* subgenus, diverged from a common ancestor ~ 27 Mya, encode both ubiquitously and germinal expressed NAC-α as well as ubiquitous NAC-β genes, while the gNAC-β genes are found in the *melanogaster* group genomes, but not in *obscura* group. The putative ancestors of gNAC-β, gNAC-β-like genes, were found in earlier branched off species, *D.willistoni* and *D.busckii*, as well as in mosquitoes, *Aedes aegypti* and *Culex quinquefasciatus*, which indicates a rather early origin of this gene in insects.

**Figure 3.**
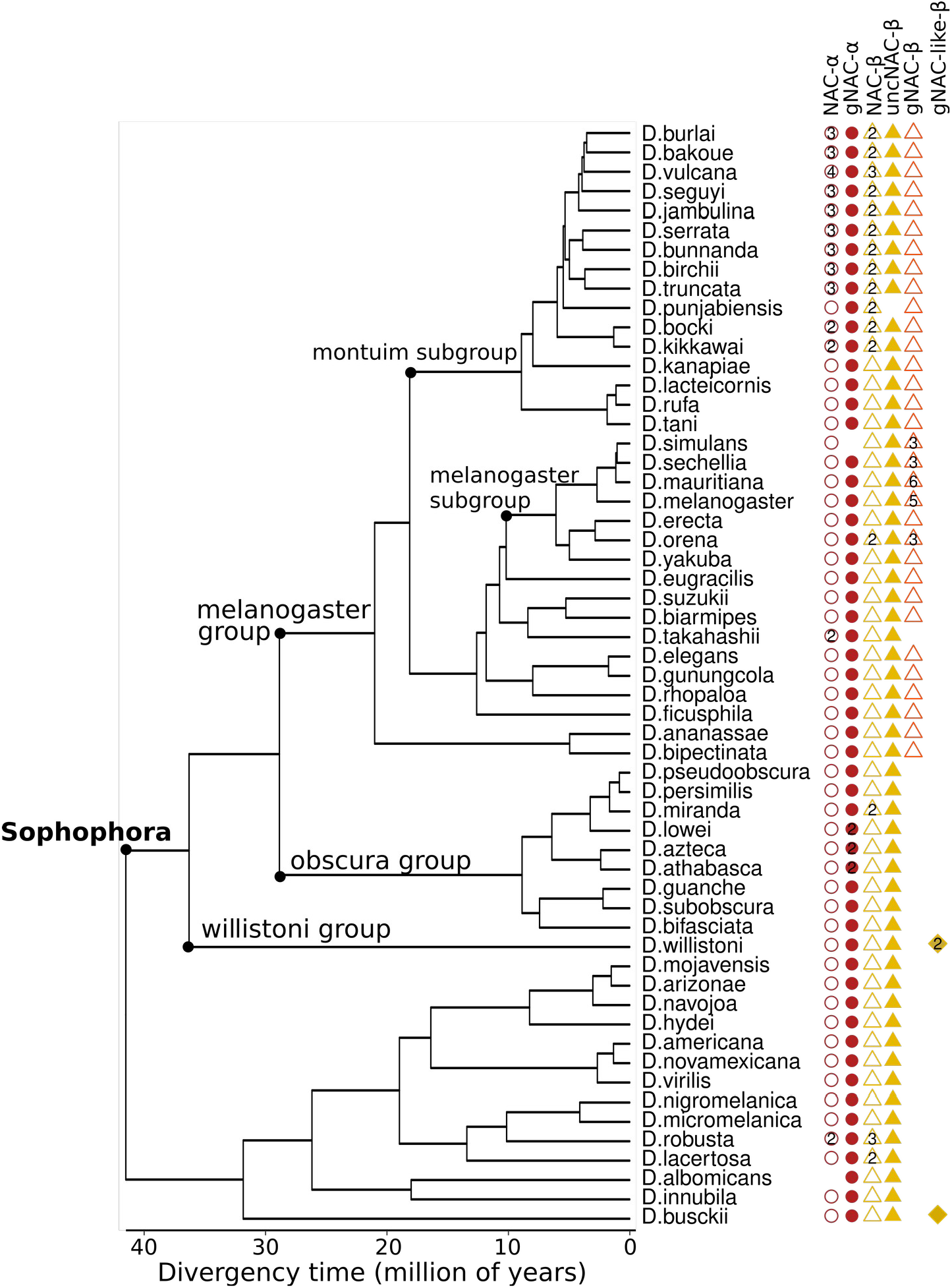
Phylogenetic tree of the Drosophila genus. The lack on this tree of some species (*mayri, watanabei, leontia, asahinai, pectinifera, obscura,montana, malanica and grimshawi*) is rather the result of the unfinished assembly of corresponding genomes preventing detection the NAC domain sequences. The number of paralogs is shown within shapes.

The numbers of ubiquitous NAC-α and NAC-β copies are shown to be correlated with each other in most species belonging to the *montium* subgroup of the *melanogaster* group. This observation is consistent with the earlier report on the production of equimolecular α- and β-subunits in yeast cells [39]. It should be noted that the correspondence between the number of genes of the germinal α- and β-subunits is lost in several species from the *melanogaster* group, which are characterized by the amplification of the gNAC-β genes.

All species, possibly with the exception of one, *D.punjabiensis* (Fig.3), contain a single copy of the uncharacterized gene *uncNAC*-β (its orthologous *melanogaster* gene structure is presented in Fig.2) encoding the NAC domain sequence followed by about 800 amino acid residues representing an extended IDR. Perhaps, its heterodimerization with the alien NAC-α domains allows to transfer the functionally important function of flexible long IDR to newly formed heterodimers. The gNAC-β-like gene (this g-designation is given only due to the greatest similarity of its NAC domain with the gNAC-β domain) is detected only in *D. willistoni* and *D. busckii* species diverged just at the root of the *Sophophora* divergence.

The cross-species alignment (Fig. S2, Data Set 1) shows that the ubiquitous NAC-α domains are very conserved, while the gNAC-α domains are diverged slightly faster (the average amino acid identity per column for multiple alignment of NAC domains is 80% and 75.6% for the ubiquitous and gNAC-α, respectively). The same feature is even more noticeable for the orthologous pairs of the NAC-β ubiquitous and germinal counterparts (the average identity per column in the multiple alignment of NAC domains is 69.4% and 38.8% for ubiquitous and germline NAC-β domains, respectively). These values for the multiple alignments of the overall IDR sequences for gNAC-β is 8%, while the distinct regions within IDR reach the average identity of 50%, indicating significant conservation. Off note, some species subgroups from *Sophophora* subgenus have several highly similar paralogs of the αNAC and/or βNAC subfamilies.

It is known that IDRs are conserved across orthologs in vast majority of cases [40]. The cross-species alignments revealed more discrepancies in the disordered regions of the gNAC subunits compared to ubiquitous ones, while these regions also contain many conserved amino acid regions (Fig.S2). The more elongated orthologous indels were detected carrying several predicted phosphosites as is shown for the gNAC-β sequences of the *melanogaster* group species, including *D. simulans*, the closely related species D. *ananassae* and *D. bipectinate* and for the gNAC-α in the *D. ficusphila* (Fig.S1 B-D; Fig.3). Here we can see that phosphorylation increases the density of a negatively charged cluster enriched in Asp/Glu (D/E) residues, which is believed to promote phase separation of disordered regions of ribosome associated protein with subsequent regulation of translation [41].

### Ectopic expression of the germinal NAC-β rescues lethal effect of the loss of its ubiquitously expressed counterpart

The question arises about the possible functional interchangeability of ubiquitous and germinal NAC subunits, in particular, the ability of the ubiquitous NAC-α-gNAC-β chimeric heterodimer to provide at least some of the diverse functions performed by ubiquitous NAC. We evaluated the ability of the expression of gNAC-β subunit to suppress the lethal mutation of the *bic* gene [13] which encodes the ubiquitously expressed NAC-β subunit.

The generated genomic insertions of transgene copies carrying gNAC-β under the actin5C promoter were tested for their abilities to suppress the lethal effect of the null *bic^1^* gene mutation [13] (Fig. 4). Transgene insertions into chromosomes 2 (T2) and 3 (T3) were shown to provide high and medium levels of gNAC-β expression, respectively, as assessed by WB analysis (Fig 4A). T3 showed no visible suppressive effect (Fig 4 B), while T2 ensured the survival of up to 30% of fertile individuals. We hypothesize that gNAC-β protein, which carries the additionally acquired C-end IDR sequence, is able of performing some functions of the ubiquitously expressed NAC-β subunit in the NAC-αβ heterodimer.

**Figure 4.**
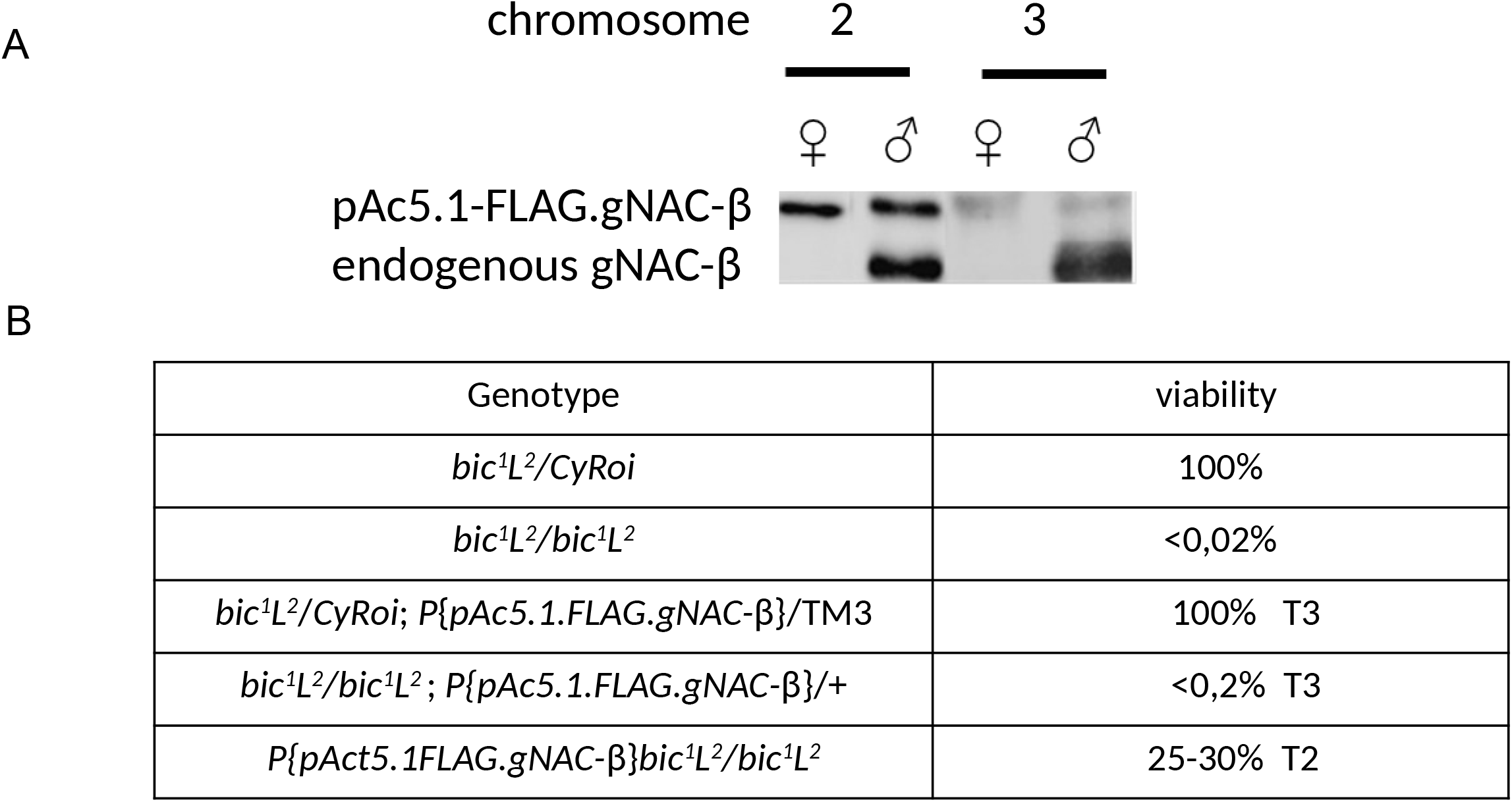
The lethal phenotype of the *bic1* mutation causing the loss of the ubiquitously expressed NAC-β is rescued by ectopic gNAC-β expression. **A.** Western blot analysis of FLAGgNAC-β transgenes (T2 and T3) expression inserted to chromosome 2 or 3, respectively. **B.** relative viability of the genotypes demonstrating transgene T2 induced suppression effect.

## Discussion

We have detected earlier the germinal testes expressed β-subunit (βNACtes) of heterodimeric NAC-αβ in *D.melanogaster* [7]. As far as we know, this was the first indication of the presence of tissue specific NAC in multicellular eukaryotes. Here we discovered the germinal NAC-αβ paralog, α-subunit of which is expressed exclusively in the germline during early embryonic development. In this paper, we also found the expression of germinal specific NAC-α and NACβ-subunits in germ cells during early embryonic development of *D.melanogaster*, which allows us to use the designation gNAC for germinal specific heterodimeric NACs. We failed earlier to observe the fertile negative effect of βNACtes siRNA KD in testes, possibly due to uncomplete depletion of the expression of these amplified genes [7], but the identification of a unique gene for the NAC-α subunit, presented in this paper, opens the way for the elimination of gNAC protein activity using CRISPR/Cas9 gene editing to trace the putative negative fertile effect.

Comparison of the amino acid sequence patterns of the ubiquitous and germinal NAC subunits revealed in gNAC significantly extended IDRs at the C- and N-ends of the β and α subunits, respectively. The burst of the interest to IDRs is explained by a large amount of recently accumulated data demonstrating the functional role of IDR, their influence on the mechanisms of protein-protein interaction and the formation of biocondensates. It is known that IDRs are protein unfolded parts that provide protein-protein interactions [42] that are considered to be necessary to perform ubiquitous NAC function and its chaperone activity in *C.elegans* [36], and we suggest germinal specific evolutionary acquired IDR sequences to participate in the sophisticated germline protein specific interactions. We also found the excess of predicted phosphorylation sites (phosphosites) at the extended IDRs of gNAC subunits compared to the ubiquitous counterparts. Taking into account the important role of NAC with its flexible adaptor sites ensuring interactions between the nascent polypeptide and components of the protein sorting machinery [1, 2] and functional conformational mobility of phosphorylated IDRs [41, 43], the in-depth experimental studies of NAC phosphorylation will be necessary to discern the peculiarities of their functions in somatic and germinal cells. However it remains a challenge to associate these IDR regions with specific biological and biochemical functions based on their amino acid sequences and putative PTM sites [44].

Here we found that the lethal effect of the lack of ubiquitous NAC-β subunit can be rescued by the ectopic extensive expression of its germinal paralog carrying evolutionary acquired extended terminal IDR sequences. This observation cannot in any way be interpreted in favor of a successful replacing the ubiquitous heterodimer with a germinal one, since the effectiveness of the chimeric gNACβ-NAC-α heterodimer in the implementation of protein homeostasis has not been evaluated. The observation of accomplished suppression indicates only a certain level of functional flexibility of the heterodimer subunits.

The detection of specific NAC in the germline is of particular interest, given some peculiarities of protein synthesis regulation in the germline development, associated with proliferation and differentiation programs programs [45]. We suggest the germinal NAC subunits, carrying the extended flexible and easily accessible for PTM amino acid sequences can be used in the germline proteostasis regulation. The recent *in vitro* study of human βNAC function using reticulocyte rabbit ribosomes has revealed a novel NAC function as a component of the ternary complex «NAC-ribosome nascent chain complex-SRP with SR-receptor» located at the ribosomal exit site [46]. The authors found ubiquitous NAC to perform selective biasing the flexible conformational landscape of multicomponent SRP for association with its receptor during the cotranslational targeting the N-terminal signal carrying nascent polypeptide to ER. Thus, a more active and broader role of NAC in proteosynthesis as an allosteric regulator of SRP functions has been demonstrated, while the possible role of the disordered region of ubiquitous β-NAC in its N-terminal region associated with the ribosome has not been mentioned [36].

The switch-like decisions during germ cell development are proposed to be closely related to the formation of large protein complexes, including large granules, also known as membraneless organelles (for review [47, 48]). In this regard, new ideas about the cotranslational assembly of multisubunit protein complexes with the participation of a chaperone, based on rigorous experiments, are of particular interest [2, 49–51]. The authors propose that the translations of subunit complexes are spatially confined. These ideas are discussed in view of compartmentalized translation principles with regulatory implications in development [52]. It would be attractive to extend such concepts to the description the complex processes of granule formation in the germ plasm compartment of a mature oocyte that can be accompanied by the gNAC in *D. melanogaster* based on our preliminary experiments. However, today, the identification of gNAC only opens the door to the first experimental approaches to explore its functions.

## Methods

### Fly stocks and crosses

*Batumi* line laboratory stock was used for immunostaining.

To generate recombinant chromosome 2 carrying the *bic ^1^* lethal allele and FLAG.gNACβ transgene, the *y^1^w^*^; Eney-whited700-pAc5.1.FLAG.gNAC*-β.*RetMI07200/CyRoi* females were crossed to *bic^1^L^2^/CyRoi* males and the F1 *Eney-whited700-pAc5.1.FLAG. gNAC-β.RetMI07200/bic^1^L^2^* females were crossed to *yw^67c23^*, +/+ males. Male progeny of this cross with recombinant chromosome (marked *whited700* and *L^2^*) *Eney-whited700-pAc5.1.FLAG.gNAC*-β.*RetMI07200 bic^1^L^2^*/+ were collected and individually crossed to *yw^67c23^*, +/*CyRoi* females. Males *Eney-whited700-pAc5.1.FLAG.gNAC*-β.*RetMI07200 bic^1^L^2^/CyRoi* were crossed to *bic^1^L^2^/CyRoi* females and the viable *Eney-whited700-pAc5.1.FLAG.gNAC*-β.*RetMI07200 bic^1^L^2^/bic^1^L^2^* individuals (lacking *CyRoi* and carrying the *whited700* and *L* markers) were traced/selected to evaluate the suppression effect of the *bic*^1^ lethality.

### Construction of plasmids expressing FLAG.gNAC-β and HA.gNAC-α proteins

RT-PCR products of gNAC-β and gNAC-α ORFs were inserted into pAc5.1.FLAG or pAc5.1.HA carrying plasmids. Oligos for plasmid insertions:

pAc5.1.FLAG.gNAC-β (CG18313)

Xho1-5’tataCTCGAGACAATGGATTTCAACAAGCGACAG

Apa1-5’ tataGGGCCCCTAATCTTCGTCCTCGGAGACCT;

pAc5.1.HA.gNAC-α (CG4415):

Kpn1-5’tataGGTACCTTCCTCAAGATGGGTAAGAAGCAGA

Xho1-5’ tataCTCGAGGTTGTCGTTCTTCAGCAGCGC

### Generation of transgenic drosophila lines expressing FLAG.gNACβ protein under pAc5.1 promoter

Transgenic strains carrying construct attBs-Eney-whited700-pAc5.1.FLAG.gNAC-β-attBsrev-pSK=aeca were generated by phiC31-mediated site-specific integration at the MiMIC site [53–55] in the *Ret* gene of chromosome 2 (Bloomington #43099, y1w*; MiRetMI07200/SM6a) and at the site (Bloomington #24862, yM{RFP[3xP3.PB] GFP[E.3xP3]=vas-int.Dm}ZH-2A w[*]; PBac{y[+]-attP-9A}VK00005) of chromosome 3. The vas-dPhiC31 strain bearing the phiC31 gene under the control of the *vasa* gene promoter on the X chromosome was used as an integrase source [54]. The germ-line transformation of the embryos was performed according to standard protocol [56] with the approximately 40% efficiency of integration.

### Cell culture transfection

Transient transfections of S2 cells were performed with the help of FuGENE^®^ HD Transfection Reagent (Promega# E2311) according to the manufacturer’s instructions. After 3-4 days after transfection cells were harvested and subjected to immunostaining and western blot analysis.

### Generation of germinal anti-NACβ antibodies

Rabbit was immunized by His-tagged germinal NAC-β subunit as described earlier [7]. Antibodies were purified by antigen affinity chromatography using Thermo Scientific AminoLink Plus Coupling Resin according to the manufacturer’s protocol (Thermo Fisher Scientific). The generated antiserum was shown to recognize exclusively the germ cells in testes and early embryos and was used in Western-blot analysis (dilution 1:1000).

### Western-blot

WB analysis was performed using rabbit polyclonal antibody to germinal NACβ, monoclonal mouse antiFLAG M2 (Sigma F3165) and Monoclonal anti HA-Tag antibodies (mAB#2367 Cell Signalling Tech) (1:1000 dilution) according to_[7].

### Immunostaining embryos

Embryos were collected at age 12-15 hour, dechorionated in bleach (2%) for 3 minutes and after thorough rinsing with water were placed in a 1:1 heptane/methanol mixture (−20°C) in a 1.5 ml microtube and devitellinized by gently shaking the tube. Devitellinized eggs, that sink in the methanol phase, were then removed and rinsed 3 times with cold methanol. At this stage eggs can be stored at −20°C until immunostaining. For immunostaining eggs were gradually rehydrated with methanol-PBT (PBS with 0,1% Tween20), washed 3 times with PBT-X (PBT with 0,3%Triton X100) and permeabilized in PBTX with 0,3% Sodium deoxycholate (Sigma) for one hour. Then embryos were washed three times PBTX and blocked with PBTX containing 5% normal goat serum (NGS, Invitrogen) for 1 hour.

Embryos then were incubated first in specific primary antibodies in PBTX containing 3% NGS overnight at +4°C and after washing 4 times in PBTX at room temperature, incubated the next overnight at +4°C with secondary antibodies labeled with Alexa in a dark chamber. Embryos were mounted in Invitrogen SlowFade Gold Antifade reagent. The following primary antibodies were used: rabbit polyclonal anti-gNACβ (1:500), rat anti-VASA (1:200) (DSGB: AB_760351). Secondary antibodies (1:500) (Invitrogen, Thermo Fisher Scientific) were the following: anti-rabbit IgG Alexa Fluor 546; anti-rat IgG Alexa Fluor 488. Confocal microscopy was done using LSM 510 META system (Zeiss).

### Phylogeny of 69 Drosophila species

To estimate the phylogeny of Drosophila species, the 192 RefSeq and GenBank genomic assemblies of 70 species were downloaded from the NCBI. Each assembly was subjected to BUSCO v.4.1.4 [38] analysis to identify the unversal single-copy orthologs from OrthoDB (-l dipteria_odb10). The genomic assembly of *Musca domestica* (GCF_000371365.1_Musca_domestica-2.0.2) was also analyzed by BUSCO for further usage as the outgroup. 3,119 universal single-copy BUSCO orthologs that are present in at least 90% of 192 assemblies and 69 genomic assemblies (one assembly per species) having at least 90% of 3,119 single-copy orthologs were selected for the *Drosophilidae* phylogenetic analysis and the identification of NAC family proteins (Table S1). The concatenated multiple sequence alignments of the orthologous proteins (using MAFFT v7.471 [57] followed by alignment trimming with trimAl v.1.4 [58] (-gt 0.5) resulted in 710,094 amino acid columns that were used to estimate the maximum likelihood species phylogeny using RAxML v.8.0 [59] with the PROTGAMMAJTT model, rooted with the *Musca domestica*. We then used r8s [60] to estimate branch lengths in terms of millions of years with four calibration points [61] 25-30 million of years (moy) for the common ancestor of *D. pseudoobscura* and *D. melanogaster*, 40 moy for the common ancestor of *D. virilis* and *D. melanogaster*, 6-15 moy for the common ancestor of *D. yakuba* and *D. melanogaster*, and 100 moy for the common ancestor of *D. melanogaster* and *Musca domestica*.

### The identification of NAC family proteins

The initial identification of NAC genes in Drosophilidae genomic assemblies was performed with *tblastn* v. 2.6.0 [62] (-evalue 10E-5) with using NAC proteins from *D. melanogaster* as the queries. The hit regions extended with the additional 2000 bp on both sides were extracted from the genomes and were subjected to the determination of the open reading frames by AUGUSTUS v. 3.1.3 [63]. The identified open reading frames were confirmed as encoding NAC domains using InterProScan v.5.39 [64] (IPR016641 for αNAC and IPR039370 for βNAC). If the analyzed genome was already annotated by the NCBI (40 genomic assemblies), then the accession number of the proteins was determined by *blastp* against the corresponding proteoms retrieved from the NCBI Protein database; otherwise (29 assemblies) the NAC protein was marked as ‘novel’. The multiple alignments of NAC proteins were carried out by the MAFFT v. 7.471 [57]. The NAC domains of αNAC and βNAC were cut from the alignments and the positions including greater than >=0.5 gaps were removed by trimAl, v.1.4 [58]. Phylogenetic analysis of NAC domains was performed using the FastTree program [65] with default parameters, with the WAG evolutionary model and the discrete gamma model with 20 rate categories. The tree structure was validated with bootstrap analysis (n=100).

## Supporting Information

**Figure S1. Predicted phosphosites in IDRs of the gNAC subunits.** Docking site for various Ser-Thr protein kinases (CK1, CK2, NEK2, LATS, ProDKin, PIK and GSK3-Ser) are indicated; X-linked paralogs of *D.melanogaster* (see Flybase.org) (A). Fragments of orthologous alignments indicate divergencies caused by indels carrying phospho-sites for gNAC-α (B) and gNAC-β (C, D).

**Figure S2.** The phylogenetic trees for the NAC-α **(A)** and NAC-β **(B)** proteins of the *Drosophilidae* genomes. The species, accession number of protein and genomic coordinates of the gene are shown. The identified protein is marked as a “novel” if its accession number is non-available at the non-annotated genome. The blue integers indicate the bootstrap values (only values greater than 0.5 are shown).

**Table S1**. The genomic assemblies of Drosophilidae species that were used for the BUSCO analysis. For each genomic assembly the number and percentage of common unique singlecopy orthologs are given. Only 69 genomic assemblies with >90% common unique singlecopy orthologs were used for the phylogenetic analysis and the identification of NAC encoding genes.

**Data Set S1.** The multiple alignments of NAC-α, gNAC-α, NAC-β, gNAC-β and uncNAC-β proteins encoding in Drosophilidae genomes. For each protein the species, the accession number of protein and genomic coordinates of the gene is shown. If the accession number of protein is not available for the non-annotated genomic assembly, then the protein is marked as “novel”.

## Acknowledgments

We thank. O. Maksimenko for her help and advice in generation rabbit antibodies to the gNAC-β protein and V. Velkov for his input in this manuscript. This work was supported by a research grant from the Russian Science Foundation (RSF) (grant number 19-74-20178).

